# The Human Organ Atlas

**DOI:** 10.1101/2025.07.31.667856

**Authors:** Claire L. Walsh, Joseph Brunet, David Stansby, Guillaume Gaisné, Yang Zhou, Maximilian Ackermann, Alexandre Bellier, Camille Berruyer, Axel Bocciarelli, Marjolaine Bodin, Bernadette S. de Bakker, Hector Dejea, Alejandro De Maria Antolinos, Klaus Engel, Andy Götz, Joseph Jacob, Daniel Jonigk, Joanna Purzycka, Theresa Urban, Stijn E. Verleden, Ruikang Xue, Paul Tafforeau, Peter D. Lee

**Affiliations:** Department of Mechanical Engineering, UCL, London, United Kingdom; ESRF, The European Synchrotron, 71 Avenue des Martyrs, CS40220, 38043 Grenoble Cedex 9, France; Centre for Advanced Research Computing, UCL, London, United Kingdom; Institute of Functional and Clinical Anatomy, University Medical Center of the Johannes Gutenberg-University Mainz, Mainz, Germany; University Grenoble Alpes, Department of Anatomy (LADAF), CIC INSERM 1406, Grenoble, France; Amsterdam UMC location University of Amsterdam, Department of Obstetrics and Gynecology, Meibergdreef 9, Amsterdam, The Netherlands; Amsterdam Reproduction and Development research institute, Amsterdam, The Netherlands; Siemens Healthineers AG Erlangen 91052, Germany; UCL Respiratory, UCL, London, United Kingdom; Satsuma Lab, Hawkes Institute, UCL, London, United Kingdom; Institute of Pathology, University Medical Center RWTH University of Aachen, Aachen, Germany; Biomedical Research in Endstage and Obstructive Lung Disease Hannover (BREATH), German Center for Lung Research (DZL), Hannover, Germany; Antwerp Surgical Training, Anatomy and Research Centre (ASTARC), University of Antwerp, Wilrijkn, Belgium

## Abstract

We present the Human Organ Atlas (HOA), an open data repository making accessible multiscale 3D imaging of human organs. The repository also provides software tools and training resources enabling worldwide access, sharing, and analysis of these datasets, facilitating further research and the continued expansion of the HOA. The images are generated using a synchrotron imaging technique - Hierarchical Phase-Contrast Tomography (HiP-CT) that uses the ESRF’s Extremely Brilliant Source, spanning whole organ imaging at around 20 μm/voxel with local volumes of interest within the intact organs imaged down to ∼ 1 μm/voxel. This offers a comprehensive exploration of human anatomy, providing unparalleled insights into intricate structures and spatial relationships. The Human Organ Atlas offers researchers, clinicians, and educators a valuable resource for anatomical study, image analysis, medical education, and large-scale data mining.

## Introduction

Human organs are intricate 3D structures. Their hierarchical organization, from the extracellular matrix to cells to functional units to tissues to organs, underpins functional properties allowing both coordination and specialization. Mapping the 3D morphology and spatial distribution of biological structures at these multiple scales can provide significant healthcare and biological impacts including bounding what constitutes healthy and pathological variability of structures, linking between morphology and transcriptome, and providing data to explicitly simulate function from structure. Efforts to map the entire human body at the cellular level and tissue level have been ongoing for some time, with 3D imaging as a cornerstone in many of these efforts [1, 2, 3, 4, 5, 6, 7]. In recent years we have developed Hierarchical Phase-Contrast Tomography (HiP-CT), a synchrotron X-ray tomography technique which creates hierarchical image volumes of ex vivo intact human organs, spanning the scales from the single cell to whole organ [8, 9]. HiP-CT imaging of a human organ has typically involved imaging at multiple resolutions: initial imaging of the whole sample with an overview scan at an isotropic voxel size of ∼20 μm, followed by zoom scans in selected volumes of interest at a configurable isotropic voxel size of ∼1 - 6 μm, all within the intact organ. The current lowest achieved voxel size is 0.65 μm. The images at each resolution can be rigidly aligned to one another creating hierarchical datasets that provide histological level detail in 3D at user selected locations within whole organs without physical sectioning. With technical developments since the original HiP-CT results, the speed and consistency of imaging have dramatically increased, allowing for routine and rapid (2 – 8 hours depending on organ size and scan parameters – see Figure 7) multi-resolution scanning of human organs which has generated a large database of 3D hierarchical organ images. HiP-CT has already shown impact in biomedical fields [10, 11, 12, 13, 14, 15, 16], but the technique requires the small x-ray source size, high energy, large beam size, and long propagation distances available on beamlines BM05 and BM18 at the European Synchrotron Radiation Facility (ESRF) [17]. This limits accessibility of the technique for those without the resource or training to independently apply it. To maximize the utility and impact of HiP-CT we have developed a data portal, The Human Organ Atlas (HOA), to ensure data we collect is findable, accessible, interoperable and reusable (FAIR) [18] (Figure 1 A). In this paper, we outline the features of the HOA (Figure 1), the organization of the data portal, and the data currently available. We then provide demonstrators for how open data in the HOA may be utilized by different communities through two examples:

1. application to machine learning segmentation utilizing multi-resolution images, and
2. application to anatomical training and education through advanced visualization techniques.

**Figure 1:**
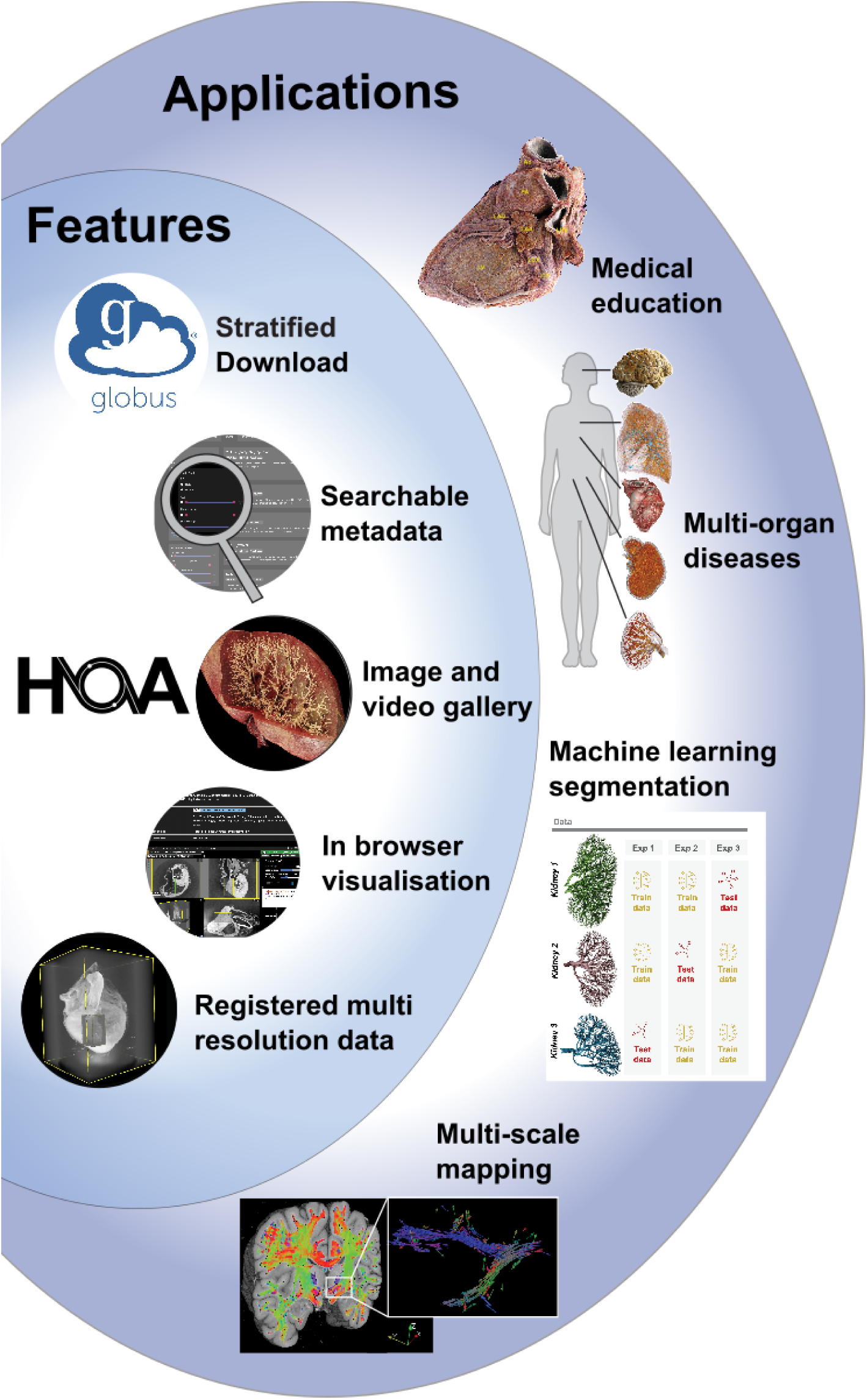
Overview of the Human Organ Atlas (HOA), demonstrating features including open data download, search tools based on metadata tags_4_, image/video galleries, browser-based visualization, and registered hierarchical datasets. “Applications” show some of possible uses that the HOA has and will enable for researchers around the world, including anatomical visualization and training, studying multisystemic disease e.g. COVID-19, hypertension etc., machine learning image segmentation challenges (panel adapted from [19]), and mapping of texture features such as white matter orientation in the brain.

We finally discuss the future direction for development of the Human Organ Atlas.

## Results

### Features and Organization of the Human Organ Atlas

The HOA data portal is available at human-organ-atlas.esrf.eu. All datasets are acquired using the HiP-CT method (Figure 2 A and B) performed on ex vivo human organs, following methods and protocols that have been previously described [8, 9]. The resulting 3D hierarchical datasets provide detailed organ overviews with multiple high resolution zoom datasets nested within (Figure 2 C).

**Figure 2:**
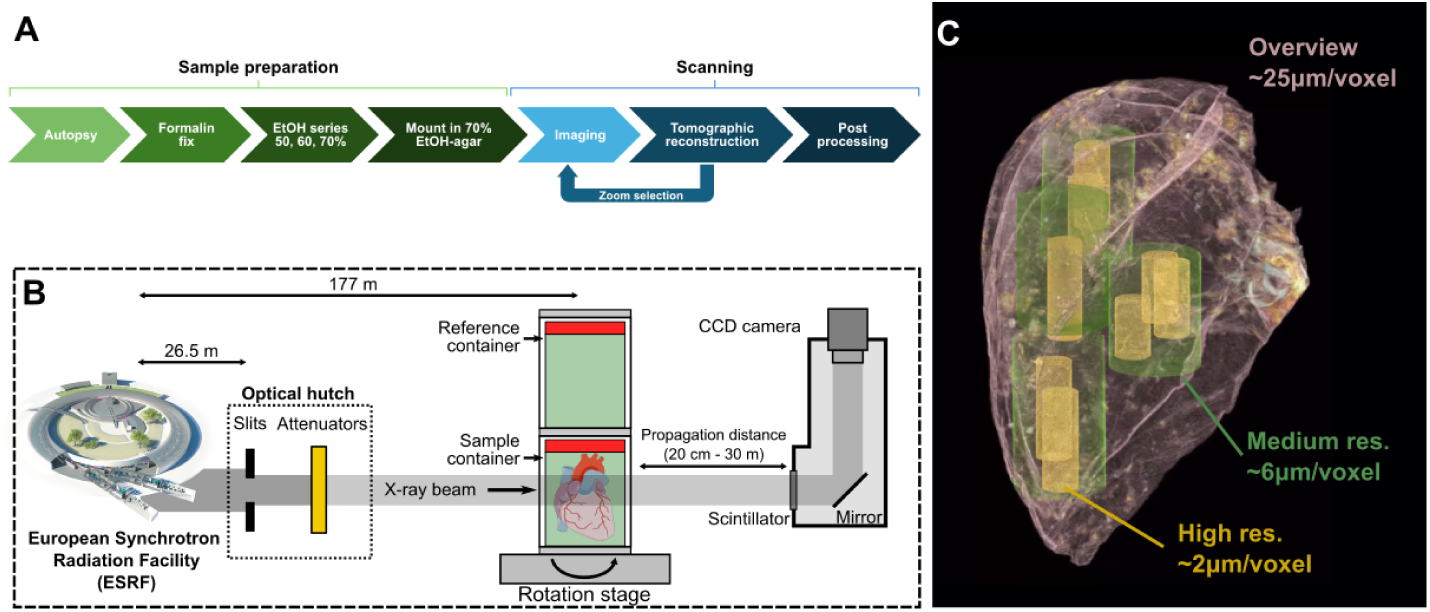
A) Overview of the HiP-CT method pipeline (adapted from [9]). B) Diagram showing the key components of the imaging setup at ESRF BM18 (adapted from [14]). C) Example of the hierarchically aligned volumes of image data from HiP-CT scan of the Lung of S-20-29. In each case there is an overview scan, and then various other resolution scans aligned within it. In this example there are medium resolution and a high-resolution scan at 6 and 2 µm/voxel respectively. Note how multiple medium and high-resolution scans can be made within a single sample, and all can be aligned to the overview scan.

The HOA portal (Figure 3 A) has features to make the data FAIR for a broad variety of users. The data are made available under a CC-BY-4.0 license which requires users of the data to cite the DOIs of the datasets they re-use.

**Figure 3:**
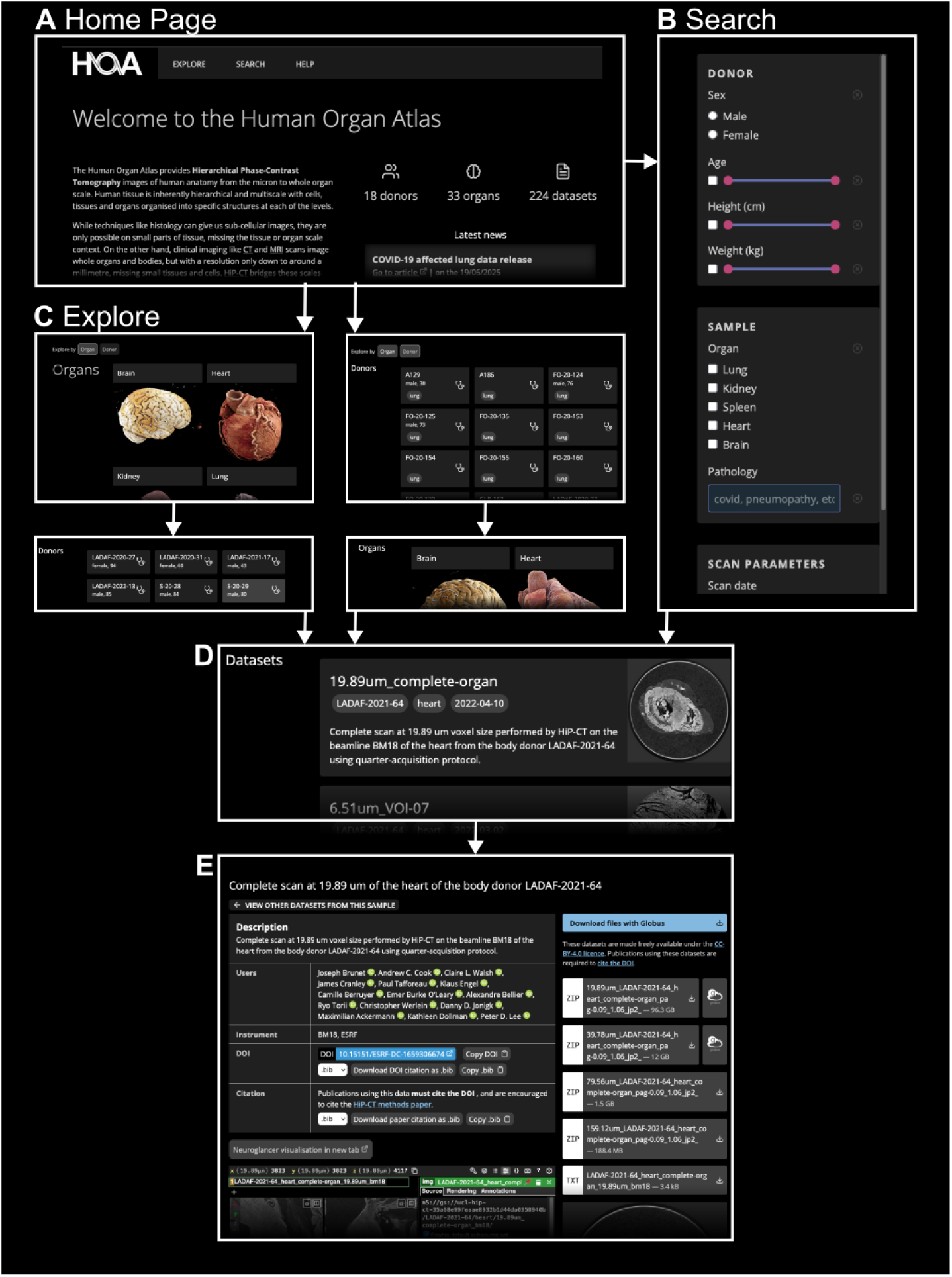
Overview of the HOA portal features. A) The HOA home page with the tabs for different pages listed across the top. To discover datasets, users can either go to B) the ‘Search’ tab to search based on medical, scan and demographic metadata., or C) the ‘Explore’ tab. Within the ‘Explore’ tab (C) datasets can be ordered first by organ (C, left column) or by donor (C, right column). From both ‘Search’ and ‘Explore’ tabs the user is then presented with D) a list of datasets, and when one dataset is chosen taken to E) the dataset page with in-browser visualisation and download options for the various down sampled datasets and metadata.

To make the data findable, the “Explore” tab allows data to be viewed either by Donor > Organ > Dataset, or Organ > Donor > Dataset (Figure 3 C); while the “Search” tab (Figure 3 B) provides a graphical interface to search and filter through the available metadata for each dataset.

To make the data accessible, each dataset page (Figure 3 E) provides data download via the Globus data transfer system (https://www.globus.org/data-transfer) (Figure 3 Bv). Data is provided and stored in the JPEG2000 image format (https://jpeg.org/jpeg2000/, [20]) with a compression factor of 10, to enable faster download and more efficient storage. Multiple down-sampled versions of the same dataset are available for download (Figure 3 E) with each dataset’s highest down-sampling level having a compressed file size of 400MB or less, making lower resolution versions of the data accessible to those with limited computational resources.

To make the data interoperable, metadata for each dataset are available for download as a JSON file that conforms to a strict schema [21]. These metadata fall under six broad categories: information about the data itself, scanning parameters, donor information, sample preparation, image registration transforms, and attribution data. A copy of the metadata for all datasets is also hosted alongside allowing for programmatic interaction with the metadata. Although best efforts are made to collect and attach as much medical history metadata as possible, in general it cannot be taken as complete for each donor due to parts of the medical history collection pipeline outside of the projects control. All the above features enable re-usability across multiple scientific and education fields, e.g. medical, bioimage analysts, medical illustrators, educators, image processing.

For each dataset a web-based visualization tool, neuroglancer [22] (Figure 3 E), allows users anywhere in the world quickly view and interact with any dataset. With just a web browser and no additional software users can assess scan quality, identify features of interest, locate zooms within each overview scan and add minimal shareable annotations before deciding whether to download the datasets for local analysis.

The higher resolution zoom datasets are registered to the appropriate whole organ overview dataset and the transforms for these registrations are provided in the dataset metadata. This powers visualisation of the zoom scans in the context of overview scans within neuroglancer on the website.

A tutorial and help page provide instruction on how to navigate and utilize the portal, as well as basic tutorials on how to download the datasets, use neuroglancer, and load and analyse data in Fiji [23]. The hoa-tools Python package [24] provides more advanced tools for data operations and processing through the scientific Python ecosystem, making use of xarray, dask, and numpy [25, 26].

### Current Data in the Human Organ Atlas

At present the HOA contains 296 3D images from 23 individual donors across 9 organs: brain, colon, heart, kidney, liver, lung, prostate, spleen, and testis (Table 1). Donors were initially sourced from two large European biobanks: The Hannover Unified Biobank (HUB) and the Laboratoire d’Anatomie Des Alpes Françaises (LADAF). Given the nature of body donation to such programs, at the time of writing the HOA is more representative of older individuals (mean age 73 years, range 30 – 94 years) and male donors with 13 male and 4 female donors. Over time the number of biobanks feeding HOA has diversified past the initial two, which will help improve the diversity of donors with future data releases. To facilitate cross-organ analysis and enable studies of multi-organ anatomical relationships, a concerted effort has been made to acquire and curate datasets for multiple organs from the same donor, notably LADAF-2021-17, for whom datasets are available for the brain, colon, heart, kidney, liver, lung, prostate, spleen, and testis (Table 1).

**Table 1:**
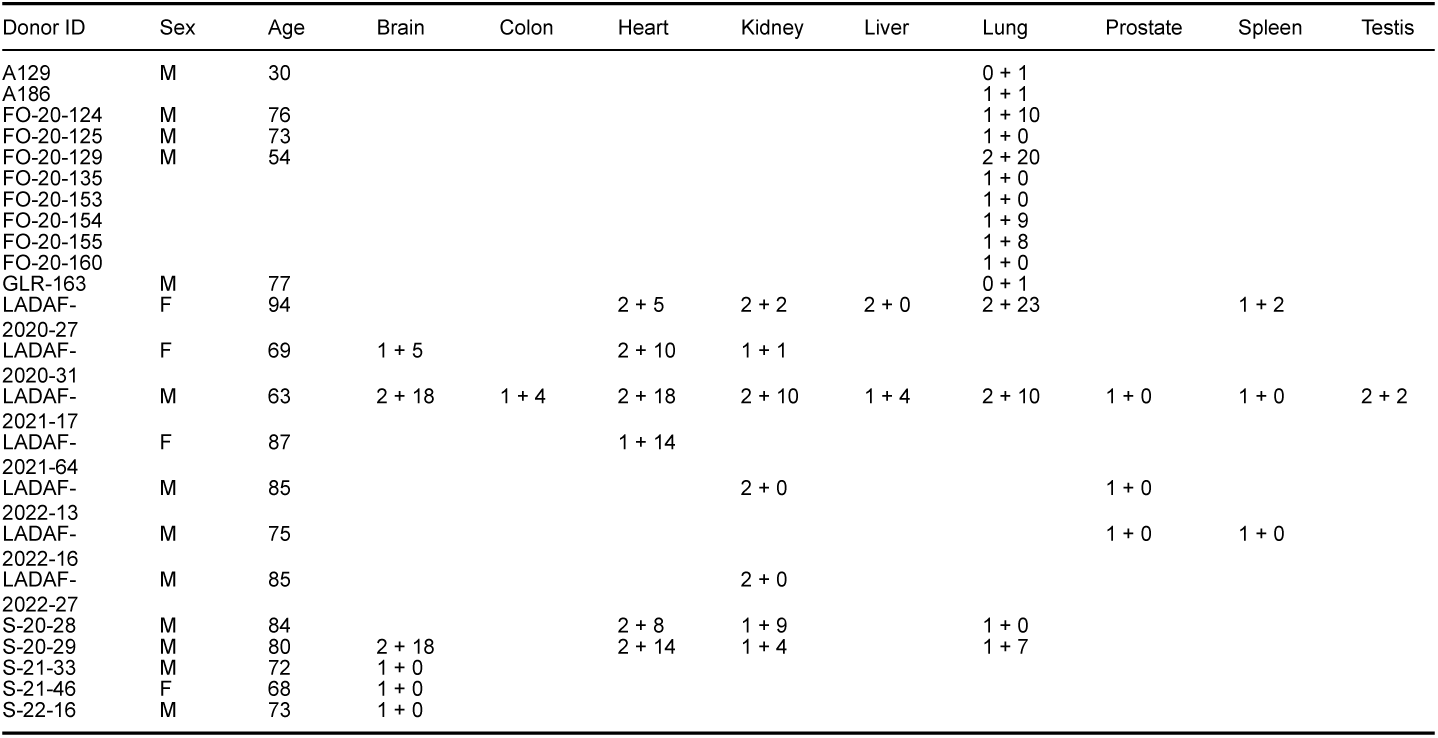
Overview of datasets available in the HOA at the time of writing. Organ columns on the right denote the number of overview datasets plus the number of zoom datasets for each organ/donor.

Within available datasets there are a large variety of total data sizes (summarized in Figure 4 A). 17 (6%) are 1TB or greater, 44 (15%) are over 500GB, and 265 (90%) are over 100GB. Dataset size does not correlate with the type of dataset (overview or zoom) and is typically in the range 30 GB – 1TB.

**Figure 4:**
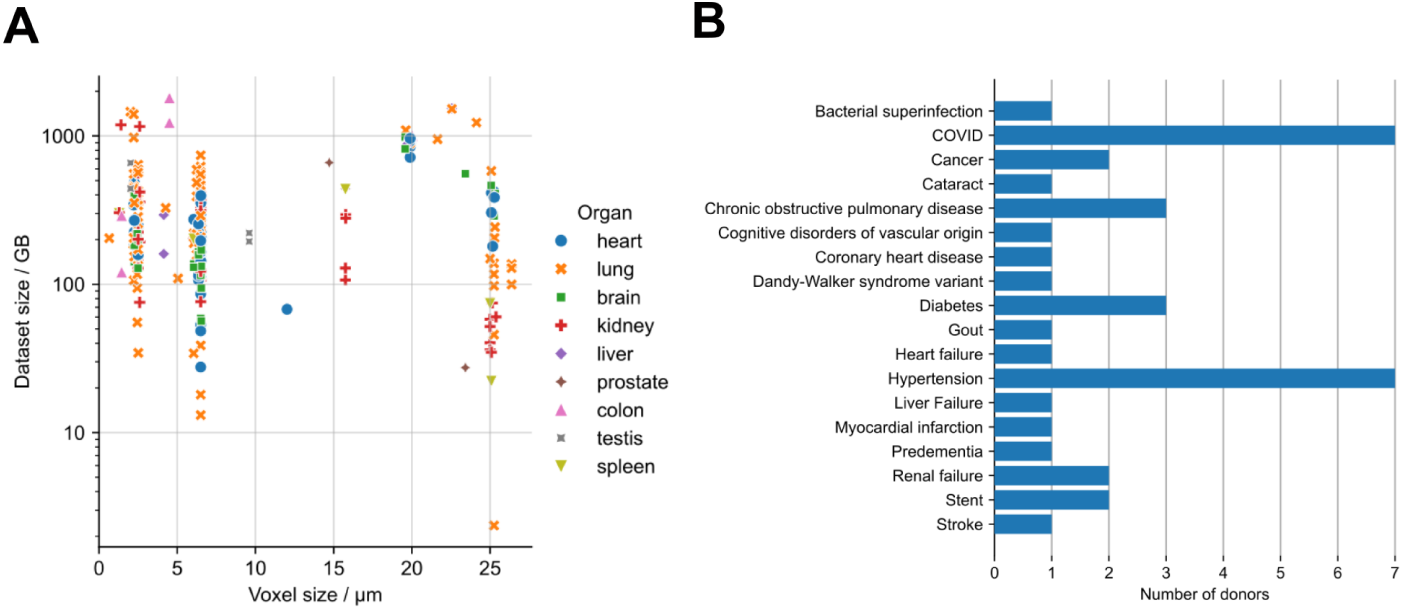
Overview of data currently in the HOA; A) the voxel size of each dataset against dataset size, with organs separated by marker style and colour. Many datasets are over 200GB and some are over 1TB, highlighting the importance of having in-browser viewing and down sampled datasets available for download. B) An overview of key medical conditions represented in the atlas. These medical conditions are provided as meta-data and allow researchers with an interest in a specific disease to find and utilise data that are relevant to their research questions. Note the higher proportion of age-related diseases such as cancer and hypertension as well as recently widespread diseases e.g. COVID-19, and some rarer diseases such as Dandy-Walker-Variant.

Given the ex vivo nature of HiP-CT and the age of donors, the HOA contains a wide variety of age-related pathologies, such as hypertension and cancer, as well as a high proportion of COVID-19 cases, reflecting both the global causes of death in recent years [27], and the roots of the HiP-CT method which was developed to better understand the changes to different organs due to COVID-19 during the pandemic. There are also rare pathologies within the HOA, e.g. in donor S-20-29 Dandy-Walker-Variant syndrome, a congenital malformation of the cerebellum which affects fewer than 1 in 30,000 individuals [28, 29] (Figure 4 B). Taken together, these samples provide a rich imaging database to study diseases that are leading causes of death in the global north today, as well as rare pathologies that few researchers may have direct access to.

### Applications in machine learning for quantification of structures

A key application of the HOA is the provision of a curated 3D hierarchical imaging dataset for the biomedical machine learning community, offering several distinct possibilities for researchers across supervised, semi-supervised, and unsupervised regimes. For example, researchers can leverage the 3D isotropic nature of the data at any one resolution to segment 3D structures, make use of the alignment of the hierarchical datasets for super-resolution applications, or make use of the large quantity of curated 3D data to train self-supervised foundation models.

For supervised approaches researchers can download datasets freely and perform their own manual segmentation of biological structures. These segmentations can then be used both directly for quantification [30, 15] or to create training datasets for supervised machine learning approaches [31, 19].

As well as supervised methods the HOA provides a large, curated 3D data pool for training unsupervised models or foundation models using e.g. masked autoencoders [32]. The limited availability of open-source largescale 3D datasets has been increasingly highlighted as a limitation for the development of 3D foundation models [33, 34]. Given the growing significance of these models including notable models such as the segment anything model (SAM) and MedSAM [35, 36, 37], and due to their capacity to provide zero-shot or one-shot segmentations [38], we anticipate this to be a large use case for HOA datasets.

A unique feature of HOA datasets that can be utilized by researchers is the hierarchical nature, i.e. aligned data of the same structures imaged at multiple resolutions. These datasets are of particular interest for super-resolution applications or machine learning based image compression [39, 40]. Another application is super-resolution segmentation i.e. propagating segmentations made on higher resolution image volumes to lower resolution image volumes within the same organ as pseudo labels (Figure 5 A). This can allow small sub-resolution structures to be mapped to whole organ imaging, enabling spatial distribution or morphology of these structures to be profiled across never examined length-scales.

**Figure 5:**
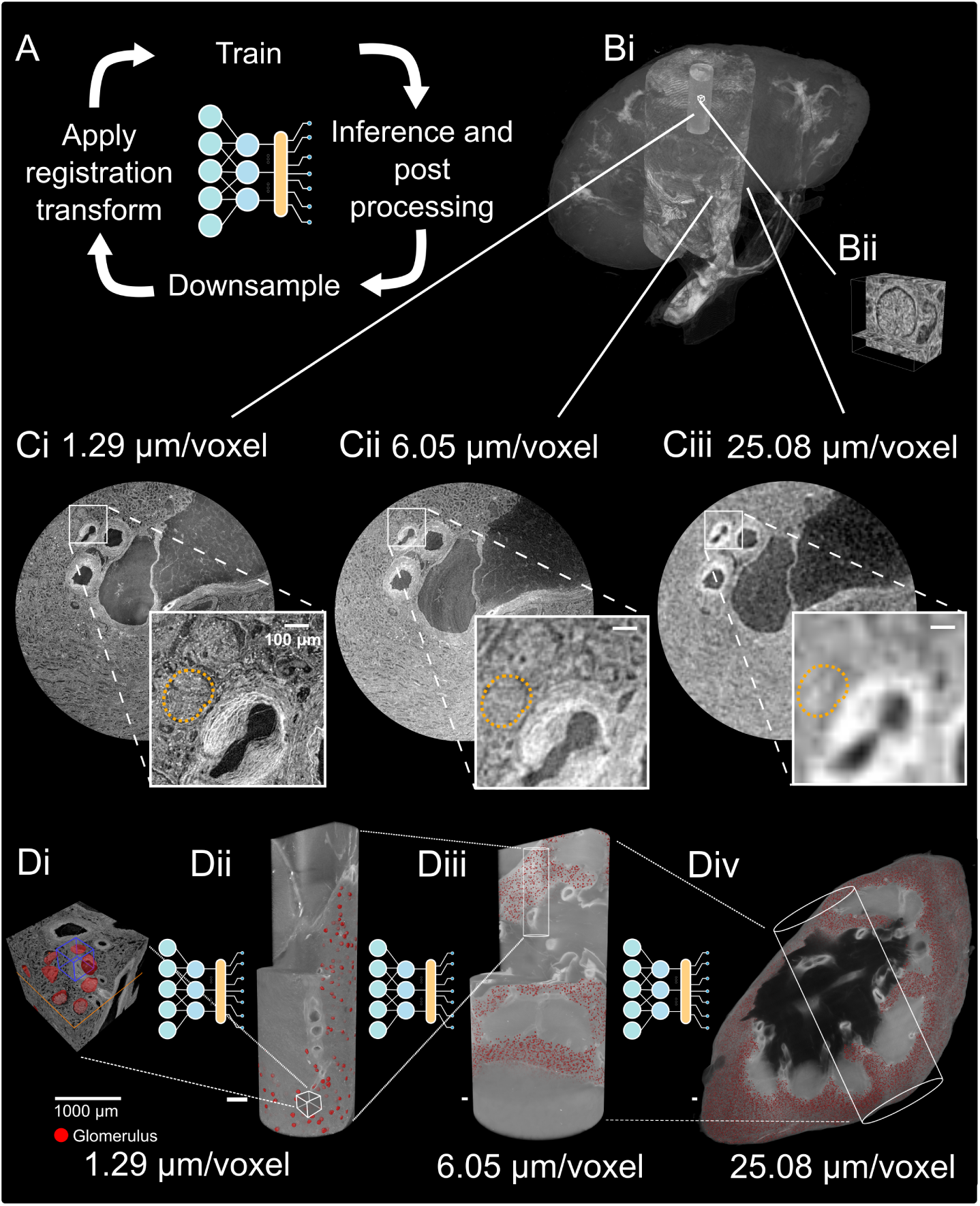
Using registered hierarchical datasets to perform a super-resolved segmentation. A) Overview of the approach. Bi) Hierarchical dataset for human left kidney LADAF-2020-27. Bii) Renal glomerulus images at 1.29 μm. Ci-iii) The glomeruli outline at each imaging resolution (orange dashed line). Note how the glomeruli cannot be reliably distinguished at 25 µm/voxel. Di-Div) The manual annotations, predictions, and inference across the resolutions with the final predicted glomeruli distribution across the whole kidney. Panel adapted from results in [41], where the multiscale segmentation pipeline implementation can be accessed at: https://github.com/UCL-MSM-Bio/2025-zhou-hipct-hierarchical-segmentation and annotated high-resolution data are at: https://doi.org/10.5281/zenodo.15397768.

As an example, in the human kidney the tissue functional unit (“the smallest tissue organization that performs a unique physiologic function and is replicated multiple times in a whole organ” [42]), is the renal glomerulus - a ball shaped tuft of capillaries approximately 100-300 μm in diameter (Figure 5 Bii). In the kidney data available for donor LADAF-2020-27, glomeruli can be easily segmented with very high accuracy in the highest resolution datasets (1.29 μm/voxel) (Figure 5 Ci), at meso-resolution (∼6 μm/voxel) glomeruli can be seen but accurate segmentation of the boundary can be challenging (Figure 5 Cii), and at the lowest resolution overview of this organ (25 μm/voxel) only very prominent glomeruli can be precisely resolved (Figure 5 Ciii).

Figure 5 Div shows the output of a multiscale segmentation pipeline applied to this kidney dataset [41], demonstrating the effectiveness of utilizing the nn-UNet model [43] as the benchmark to propagate glomeruli segmentation to the scale of the whole kidney. Such a concept is applicable to other functional units across different organs [44].

### Applications to anatomical visualization and medical education

Datasets in the HOA surpass clinical CT or ex vivo MRI [45] voxel size by one to two orders of magnitude, and unlike whole organ light sheet microscopy [46] offer isotropic voxel shape. Our voxel size is comparable to state-of-the-art whole organ serial-section histology [2], but unlike histology, carrying out the imaging using HiP-CT is non-destructive, maintaining anatomical integrity and allowing imaging all the way from the organ to the cellular scale. This and the increasing number of donors and organs available, makes the HOA a valuable tool for anatomy education and research. Visualizing detailed 3D structures, such as heart valves (Figure 6 A) and kidney vasculature (Figure 6 B) is possible at multiple scales. Unlike idealized anatomical models, real patient image data offers a more nuanced understanding of human anatomy. The application of HiP-CT for anatomy in a research context in health and disease has been demonstrated through application to the cardiac conduction system [14], hippocampal arteries of the brain [13], and 3D structure of congenital airway malformation [16]. In the medical education field, 3D modelling can be used to transfer schematic anatomical knowledge to real anatomy applicable in clinical or surgical practice.

**Figure 6:**
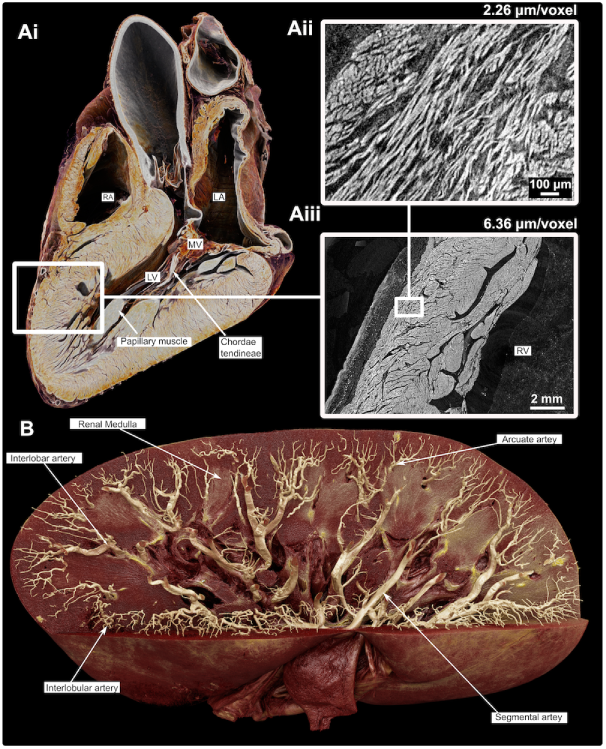
Application of HiP-CT data to creating anatomical teaching materials. A) A full human heart imaged at 19.85 µm/voxel, allowing detailed visualization of the ventricular myocardium within the intact organ. Ai displays a virtual longitudinal section with anatomical landmarks labelled, including the left (LV) and right (RV) ventricles, mitral valve (MV), papillary muscle, and chordae tendineae. Aii and Aiii show hierarchical zoom-ins of the left ventricular wall, imaged at 6.36 µm/voxel and 2.26 µm/voxel respectively, revealing the detailed myocardial microstructure such as cardiomyocyte fibres. B) Showing an intact human kidney imaged at 25 μm/voxel where the arterial tree has been segmented prior to rendering. Each anatomically defined vessel is shown in a real example from the renal artery ca. 2cm in diameter down to the interlobular arterioles ca. 50 μm radius

Combining HOA data with advanced visualization and rendering techniques, such as 3D Gaussian Splatting [47], has demonstrated the feasibility of interacting with 3D rendered HOA data, e.g. manipulating, rotating, and zooming in on specific structures in real-time. 3D Gaussian splatting significantly reduces the size of the data while maintaining high quality. This interactivity facilitates active learning, helping students understand spatial relationships between complex 3D anatomical structures, meaning that the HOA will have a central role in the next generation of anatomy education applications.

### Discussion and future directions

The Human Organ Atlas is a highly unique resource, serving many multiresolution image datasets across multiple donors, organs, and organ types; and providing metadata, online tools for visualization, and tutorials for the download and use of the datasets. This offers a comprehensive exploration of human anatomy, providing insights into the intricate structure and spatial relationships, from the whole organ to organ functional units and even down to some cellular structures. Making the data publicly available is already paying dividends, with studies outside our immediate team already making use of it [48, 49, 50, 51, 52].

For several donors, multiple organs are available in the portal (Table 1), providing an opportunity for the study of multi-systemic diseases. For example, donor LADAF-2020-27 who had a history of hypertension has lung, heart, kidney, and spleen images available. Studying the impact of different diseases across the whole body is a primary avenue of research the HOA will enable in the coming years.

The HOA continues to progress and evolve with more datasets, tutorials, and tools, and increasing data quality linked to the technical developments on the BM05 and BM18 beamlines at ESRF. Most data released so far were acquired at BM05 in the period before 2023. Since then, using the new BM18 beamline has enabled increased propagation distances and larger beam sizes, increasing the speed and contrast sensitivity of scans [17]. The HOA project is still actively collecting data and releasing these datasets to expand the diversity, quality and size of data available.

A consortium has recently been formed at the ESRF around the Human Organ Atlas and HiP-CT imaging technique - the Human Organ Atlas Hub (HOAHub, https://mecheng.ucl.ac.uk/HOAHub). The HOAHub continues to collect data and to improve upon automated analysis methods, multimodal pipelines, sample curation for specific biomedical questions, as well as dynamic image-based modelling. Through the efforts of the consortium, data will continue to be released over the coming years. Alongside the imaging datasets, we plan to start providing derived datasets that have and can be used in further analysis, for example manual segmentations.

We also plan to expand the tutorials and tools through linking to tools for simple data handling, more advanced image registration, and segmentation models, which will be provided as they are developed. Finally, we plan to make continuous improvements to the metadata, which include making it interoperable with community adopted metadata ontologies (e.g., Uberon [53]).

The HOA offers researchers, clinicians, and educators a valuable resource for anatomical study, image analysis, medical education, and large-scale data mining through machine learning. With its user-friendly interface, searchable database and ever-growing number of datasets, the HOA provides FAIR data access to a unique imaging technique. By opening the data and technique up to a wider audience, we aim for scientists across various disciplines to advance understanding, diagnosis, and treatment in the realm of human health and medicine.

## Materials and Methods

### Sample preparation and scanning

All organs were prepared following the protocols described in [8]. All the scans were performed on the beamlines BM05 and BM18 of the ESRF (European Synchrotron Radiation Facility; Grenoble, France) following the protocols of [9]. Various configurations of optics and detectors were used and are provided for each dataset in their metadata.

### Data Reconstruction and Formatting

The tomographic reconstruction process followed is detailed in [8]. After tomographic reconstruction, the 3D volumes are converted to JPEG2000 format. A lossy compression factor reduces the sizes of stored data by a factor of 10. The full resolution data are progressively down sampled by powers of 2 into lower resolution datasets to provide smaller file size options for download. This ensures the lowest resolution dataset can fit into a wider range of hardware available to researchers. The 3D volumes are then ingested to the HOA portal via ESRF servers. All datasets are also converted to the chunked N5 [54] (older datasets) or OME-Zarr [55] (newer datasets) data format and stored online as a source for the browser-based neuroglancer visualisation tool [22].

### Dataset Registration

Each zoom dataset is registered to a corresponding respective overview dataset, using software available at [56]. A common point in both datasets is manually selected to initialize the transform between the two datasets. A 128 x 128 x 32 voxel sub-volume centred on this common point is taken from the overview dataset, and the corresponding sub-volume in physical space taken from the zoom dataset. A rigid registration is then run with five parameters: the isotropic voxel scale, three translation values, and rotation around the z-axis. This registration method is designed to provide a quick and reasonably accurate registration, allowing similar structures to be easily located across zoom and overview datasets. It also provides a starting transform for users who need single voxel accurate registrations to run their own more precise registrations. The resulting transforms for each zoom dataset are provided in their metadata file.

### Metadata Collection

Metadata collation is done through a mixture of manual process and automated pipelines. Collection of donor demographic and medical data is done in accordance with ethics guidelines of the respective biobanks, and sample preparation metadata are manually recorded. Scanning and reconstruction/image volume metadata are automatically collected during the reconstruction pipeline. These metadata are collated into a single structure and validated for consistency against a schema [21].

### Website Technology

The portal is developed using Node.js and PNPM for package management. It is written using the JavaScript library React and TypeScript for maintainability and improvements over time. Vite is used to bundle the website, and it is served by a Nginx machine, deployed using Docker, on the ESRF’s servers.

The data curation and archiving within the portal leverages the DRAC (Data Repository for Advancing sCience) ecosystem [57], specifically designed to comply with the ESRF’s data policy [58]. This policy aligns with Open Science and FAIR data principles, allowing users to publish results derived from data collected at the ESRF with unique DOIs. It guarantees that metadata are archived forever and aims to retain data for at least 10 years with the option of longer periods for high-value datasets. The ESRF repository is CoreTrustSeal certified [59], underlining its adherence to internationally recognized standards for reliable and sustainable data curation. DRAC integrates key components like ICAT [60] and ICAT+ for metadata management, sample tracking and experiment logging, ensuring that all research data follows a structured lifecycle from acquisition to long-term preservation.

## Supplementary information

**Figure 7:**
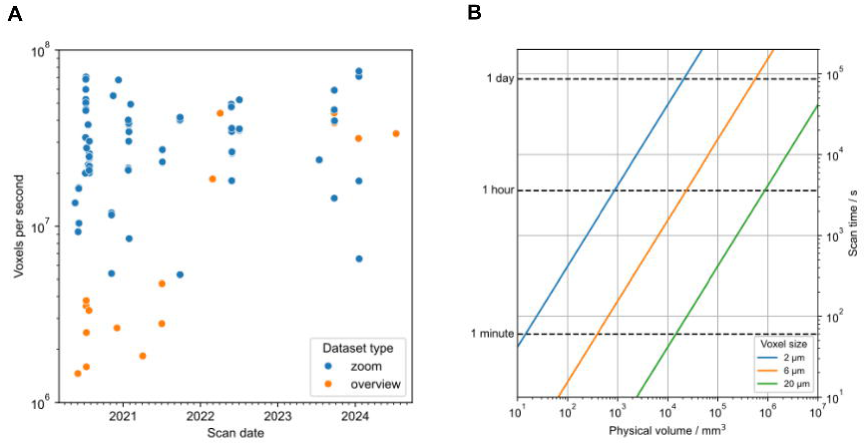
S1A shows scanning speed (voxels / second) over time for data in the Human Organ Atlas. Zoom datasets (blue points) are typically scanned at speeds between 2 x 10^7^ and 8 x 10^7^ voxels/s. Overview datasets (orange points) show an improvement in scanning speed over time due to technical developments, from around 2 x 10^6^ voxels/s earlier in the project to similar speeds as zoom datasets for more recent datasets. S1B shows scan time for a given physical volume at 2μm (blue line), 6μm (orange line), and 20μm (green line) voxel sizes, assuming a scan speed of 3 x 10^7^ voxels/second. Typical scan times for large organs at 20μm voxel size are a few hours, with sub-hour scan times at 20μm voxel size for smaller organs.

## Acknowledgments

The authors thank those who donated their bodies to science so that anatomical research could be performed. We acknowledge the European Synchrotron Radiation Facility (ESRF) for provision of synchrotron radiation facilities under proposal numbers MD-1252, MD-1290, and MD-1389. We thank Emily Newton, Alissa Parmenter, and Matthieu Chourrout for co-creating the HOA logo.

## Funding

Chan Zuckerberg Initiative grant CZIF2021-006424 (CLW, JB, DS, GG, CB, RX, PDL) Chan Zuckerberg Initiative grant 2022-316777 (CLW, JB, DS, GG, CB, HD, TU, RX, PDL) Chan Zuckerberg Initiative grant CZIF2024-009938 (JJ) The NIH BRAIN Initiative CONNECTS program via National Institute Of Neurological Disorders And Stroke (NINDS) and National Institute Of Mental Health (NIMH) award UM1-NS132358 (CLW, DS) The Wellcome Trust grant number 209553/Z/17/Z (JJ) The Wellcome Trust grant number 227835/Z/23/Z (JJ) The Wellcome Trust grant number 310796/Z/24/Z (TU) The NIHR UCLH Biomedical Research Centre (JJ) Siemens Healthineers AG (KE) The special research fund from the university of Antwerp (SEV) The Collen-Francqui Foundation (SEV) The Royal Academy of Engineering grant CiET1819/10 (PDL) The Multiscale Map of the Human Body program via a CIFAR fellowship (PDL)

## Author contributions

Conceptualization: CLW, PT, PDL Data curation: JB, DS, GG, CB, HD, JP, TU, PT Funding acquisition: CLW, PT, PDL Investigation: CLW, JB, MA, ABellier, CB, HD, JJ, DJ, JP, TU, SEV, PT, PDL Project administration: CLW, JB, MA, ABellier, BSdB, HD, AG, DJ, SEV, RX, PT, PDL Software: JB, DS, GG, ABocciarelli, MB, ADMA, AG Visualisation: CLW, JB, DS, YZ, KE Writing – original draft: CLW Writing – review & editing: CLW, JB, DS, GG, YZ, SEV, PT, PDL

## Competing interests

KE receives wages from Siemens Healthineers AG and holds stocks as well as stock options from Siemens, Siemens Energy and Siemens Healthineers. JJ declares consultancy fees from Boehringer Ingelheim, F. Hoffmann-La Roche, GlaxoSmithKline, NHSX; fees from advisory Boards for Boehringer Ingelheim, F. Hoffmann-La Roche; lecture fees from Boehringer Ingelheim, F. Hoffmann-La Roche, Takeda; grant funding from GlaxoSmithKline, Wellcome Trust, Microsoft Research, Gilead Sciences, Chan Zuckerberg Initiative and UK patent application numbers: 2113765.8 and GB2211487.0. All other authors declare they have no competing interests.

## Data and materials availability

Data used for rendering in Figure 1 is available at [61]. Code to reproduce graphs in Figure 4 is available at https://github.com/HumanOrganAtlas/hoapaper-stats. Metadata used to compile Figure 4 and Table 1 is available at [21]. Data shown in Figure 5 is available at [62, 63, 64]. Data shown in Figure 6 is available at [65, 66, 67, 68]

## Notes

https://human-organ-atlas.esrf.fr/

